# Phosphatidylinositol (4,5)-bisphosphate drives the formation of EGFR and EphA2 complexes

**DOI:** 10.1101/2024.05.03.592400

**Authors:** Pradeep Kumar Singh, Jennifer A Rybak, Ryan J Schuck, Francisco N Barrera, Adam W. Smith

## Abstract

Receptor tyrosine kinases (RTKs) regulate many cellular functions and are important targets in pharmaceutical development, particularly in cancer treatment. EGFR and EphA2 are two key RTKs that are associated with oncogenic phenotypes. Several studies have reported functional interplay between these receptors, but the mechanism of interaction is still unresolved. Here we utilize a time-resolved fluorescence spectroscopy called PIE-FCCS to resolve EGFR and EphA2 interactions in live cells. We tested the role of ligands and found that EGF, but not ephrin A1 (EA1), stimulated hetero-multimerization between the receptors. To determine the effect of anionic lipids, we targeted phospholipase C (PLC) activity to alter the abundance of phosphatidylinositol (4,5)-bisphosphate (PIP_2_). We found that higher PIP_2_ levels increased homo-multimerization of both EGFR and EphA2, as well as hetero-multimerization. This study provides a direct characterization of EGFR and EphA2 interactions in live cells and shows that PIP_2_ can have a substantial effect on the spatial organization of RTKs.

## Introduction

Cell surface proteins are crucial for responding to stimuli and communicating with surrounding cells. Understanding the molecular level interactions of proteins in the plasma membrane is vital to targeting them in drug development(1, 2). The human genome contains 58 receptor tyrosine kinases (RTKs) that share similar downstream pathways(3, 4). RTKs play crucial roles in cell growth, migration, and differentiation and are common targets of cancer therapies(5, 6). RTKs also share common structural features, including a single transmembrane helix that connects an extracellular domain to an intracellular kinase domain. In the canonical model for RTK activation, kinase function is initiated by ligand-induced dimerization(4). However, many RTKs display alternate activation mechanisms including ligand-induced conformational changes to pre-existing dimers and the formation of higher-order oligomers(4, 7, 8). Increasingly, cross-interactions between RTKs are seen as an important aspect of their function(5). Many early efforts were made to characterize heterodimers within RTK subfamilies, and there has recently been additional focus on heterodimerization across subfamilies(5, 9).

The most studied cross-family interaction is between EGFR and EphA2(5). Individually, EGFR and EphA2 each have important cellular functions. EGFR is a member of the HER (or ErbB) family of RTKs and plays a role in cell proliferation, motility, growth, and differentiation(4, 7, 10). EphA2 is a member of the largest sub-family of RTKs, the Eph receptors, and is essential to development(11-13). EphA2 is implicated in many processes that affect cellular behaviors between cells, including adhesion and repulsion(11, 13, 14). Additionally, both receptors are implicated in cancer. Although EphA2 expression is usually low in normal adult tissue, EphA2 overexpression in certain tumors is often correlated with poor prognosis(15-17). The Smith Lab and collaborators recently showed how different multimerization interfaces contributed to distinct downstream signaling and oncogenic functions(18). EGFR contributes to the development and progression of cancer and is a validated target for therapeutic intervention in various solid tumors(10, 18-22). Importantly, EphA2 has been implicated in anti-EGFR cancer therapy, where its role in drug resistance has been ascribed to direct interactions with EGFR and ErbB2(23, 24).

Because of EGFR and EphA2’s relationship in cancer, recent efforts have focused on the characterization of their functional interaction(25-28). Binding between EphA2 and EGFR is proposed to activate Ras/MAPK and RhoA signaling pathways(26, 27). However, several studies have proposed that EGFR/EphA2 heterodimers can lead to the recruitment of different adaptor proteins, and thus different biological effects(29, 30). For example, it was reported that Ephexin A1 bridges the interaction between EphA2 and EGFR(30), inducing oncogenic, noncanonical signaling through AKT pathways in the presence of EGF(26). In cases where EphA2 is overexpressed or phosphorylated at S897, there is an increased likelihood of interacting with EGFR(24, 25, 30). Despite these previous reports of EGFR-EphA2 cross-talk, there is insufficient experimental evidence to characterize the interactions and determine the factors that affect the interactions.

One factor that we expect to affect EGFR-EphA2 interactions is the membrane lipid environment, especially anionic lipids, which play an important role in RTK structure and function(31, 32). The juxtamembrane (JM) domain of all human RTKs contains several positively charged amino acid residues that interact with negatively charged lipids(33). Molecular dynamic simulations suggest that in the absence of ligands, negatively charged membrane lipids interact with and stabilize the inactive state of EGFR(34). Live cell imaging of EGFR using dSTORM, showed that the lipid PIP_2_ induces the formation of nanoclusters(35). Simulation studies suggested that EphA2 homo-multimerization is impacted by negatively charged membrane lipids, similar to the effect seen in EGFR(36, 37). Previous work from Barrera and coworkers revealed that dimerization of the isolated TM domain of EphA2 can be promoted by PIP_2_(38). However, the effect of PIP_2_ on dimerization of full-length EphA2 has not yet been investigated. Additionally, it is not known if lipids regulate the interaction between EphA2 and EGFR. In this study, we investigate the hypothesis that PIP_2_ contributes to the multimerization state of EphA2 and EGFR receptors.

Many biophysical techniques have been developed to characterize RTK multimers and their impact on cell signaling(5, 39, 40). Single-molecule (SM) fluorescence approaches are an important class of methods that have been developed to resolve membrane protein interactions in cells(40, 41). However, these SM methods are typically limited to very low expression levels (<1 molecule/μm^2^), far below the physiological expression level of many RTKs(41). Fluorescence fluctuation spectroscopy (FFS) has emerged as a valuable method for resolving membrane protein multimerization(42-45). FFS methods use time-dependent fluctuations to circumvent the optical diffraction limit. In this study, we utilize a two-color FFS technique called pulsed-interleaved excitation fluorescence cross-correlation spectroscopy (PIE-FCCS)(46) to examine the interaction between EGFR and EphA2 in cell membranes. PIE-FCCS can resolve membrane protein expression levels, diffusion coefficients, and the degree of cross-correlation (abbreviated as *f_c_*), which is a direct measure of how strongly the proteins diffuse together as dimers, trimers, and small oligomers(43). It is primarily sensitive to diffusing proteins and so does not detect immobile aggregates nor internalized proteins. As the degree of cross-correlation depends on correlated diffusion, we can interpret the co-diffusing species as stable over the timescale of the transit time (∼10^-1^ s), although it cannot resolve association lifetimes directly. The measurements described below were taken in live cells at physiological expression levels and thus provide direct evidence for membrane protein interactions in situ.

Our initial PIE-FCCS measurements did not detect interactions between EphA2 and EGFR in resting cells. This surprising result was then investigated using various ligand binding conditions and PIP_2_ modulation studies to determine the factors that affect EGFR-EphA2 interactions. By testing the effects of ligands and PIP_2_-modifying compounds, we discovered that both EGF and increased PIP_2_ levels led to significant EGFR-EphA2 hetero-multimerization. We then tested the phosphorylation levels under these conditions and found that the PIP_2_-induced multimers were less active than ligand-induced multimers, suggesting that PIP_2_ was not stabilizing an active conformation of the receptors. Overall, our work demonstrates the important role of PIP_2_ in modulating the degree of interaction between EGFR and EphA2. The results suggest that other putative RTK interactions could also be affected by PIP_2_ levels in the plasma membrane.

## Results

### EGFR does not associate with EphA2 in resting cells

To investigate the interaction between EGFR and EphA2, we performed PIE-FCCS measurements utilizing two pulsed lasers and time-correlated single photon counting (TCSPC) (Fig. 1A). We transfected cells with expression vectors encoding C-terminally tagged fluorescent proteins (eGFP and mCherry (mCH)), beginning with SRC and GCN4 as controls for monomer and dimer states, respectively(47). These control constructs have a short, lipidated peptide sequence for membrane anchoring and, in the case of GCN4, an α-helical leucine zipper motif for dimerization. We then collected data from individual COS-7 cells expressing the exogenous proteins in a physiological range of 100 to 1500 molecules/µm^2^ in the plasma membrane. From the correlation functions (Fig. 1D, E, & F), we obtained the fraction of cross-correlation (*f_c_*) and the effective diffusion coefficients of the eGFP and mCH-labeled proteins as described previously(43, 48, 49). The *f_c_*values are indicative of co-diffusion of the two receptors. The median *f_c_*value of 0.01 for SRC (Fig. 1C, light gray) and 0.15 for GCN4 (Fig. 1C, medium gray) are indicative of their monomeric and dimeric state on the plasma membrane. The dimer *f_c_* value of 0.15 is smaller than that observed for a covalent dimer (e.g. duplex DNA). The *f_c_* values from live cell measurements like those used here have been explained previously using a stochastic model(43, 47). Briefly, the cross-correlation is affected by volume mismatch in the detection system, fluorescent protein dark states, and stochastic association of red and green fluorescent labels. We next co-expressed EGFR-mCH (Fig. 1B, top) and EphA2-GFP (Fig. 1B, bottom) under ligand-free conditions. The median *f_c_* value for EGFR and EphA2 was 0.04 (Fig. 1C, dark grey), indicating that they do not hetero-multimerize in the absence of ligand. These surprising results seem to contradict earlier reports of direct interactions between EGFR and EphA2(25). In the next section we investigate the hypothesis that PIP_2_ levels in the plasma membrane could affect the degree of interaction between these two RTKs.

**Figure 1.**
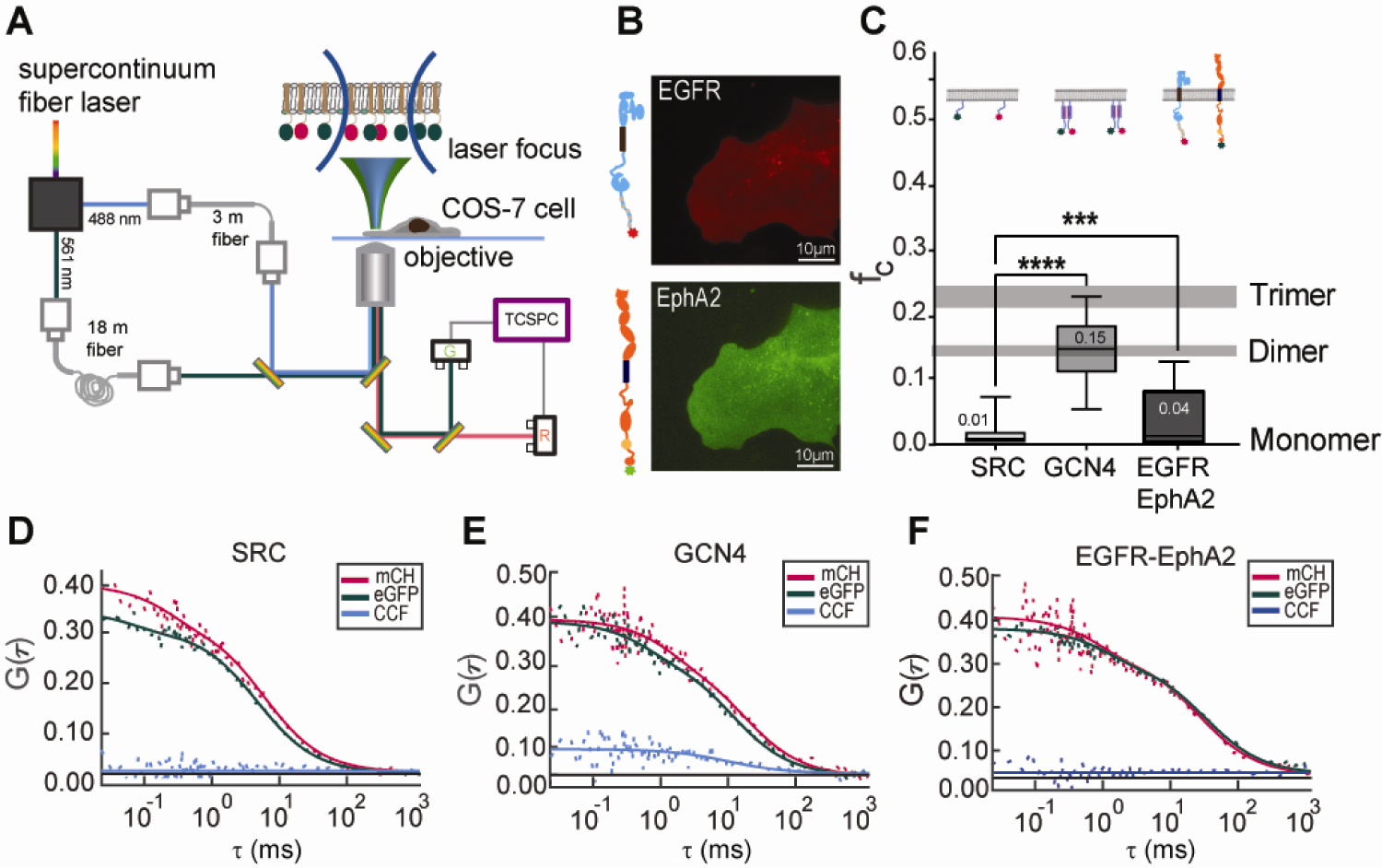
Sketch layout of PIE-FCCS and its role in membrane protien interaction. **(A)** This diagram illustrates the optical path of dual-color PIE-FCCS. A super-continuum, pulsed laser generated excitation beams (488 and 561 nm) that were directed into the sample by a dichroic mirror and microscope objective. The lasers were focused to a diameter of about 250 nm on the plasma membrane of live cells. Fluorescence emission was collected by the objective, filtered, and directed to single photon counting modules connected to a TCSPC module for data recording. **(B)** Epifluorescence images are shown for COS-7 cells expressing EGFR-mCH (top) and EphA2-eGFP (bottom). **(C)** Summary data from independent experiments collected on five different days of single cell measurements is shown for the controls and EGFR with EphA2 (*** = P<0.0004). **(D, E, and F)** Representative single-cell PIE-FCCS data is shown for a monomer control (SRC), dimer control (GCN4), and EGFR with EphA2 in COS-7 cells, respectively. In each graph, the green line is the autocorrelation function (ACF) for eGFP, the red line is the ACF for mCH, and the blue line is the cross-correlation function (CCF) between both protein constructs.

### A PLC inhibitor increases EGFR homodimerization and EphA2 homo-multimerization

PIP_2_ lipids play an important role in RTK signaling, despite constituting a minor portion of the total phospholipid composition of cell membranes(50, 51). In cell signaling, PIP_2_ is broken down by phospholipase C (PLC) into the second messengers: diacylglycerol (DAG) and inositol 1,4,5-triphosphate (IP3)(52). PLC-lll1 is expressed in most cell types and is activated downstream of RTKs(52). Several activators and inhibitors of PLC have been utilized to understand its role in cell signaling and development. U73122 is an inhibitor of PLC activity(53, 54). m-3M3FBS is a PLC activator(55, 56). In our current work, we utilize these two drug molecules to regulate PLC activity and thus increase (with U73122) or decrease (with 3M3FBS) PIP_2_ lipid density in the plasma membrane. As a control, we performed live cell imaging experiments in an assay adapted from Balla et al (57), which confirmed that U73122 increased while 3M3FBS decreased plasma membrane levels of PIP_2_.

We began by investigating how EGFR and EphA2 ligands and PIP_2_-modifying drugs affect the homo-multimerization of each receptor. For EGFR, we co-expressed EGFR-mCH and EGFR-eGFP in COS-7 cells. The PIE-FCCS measurements were recorded at expression densities between 100 and 1400 molecules/µm^2^, before and after treatment. In untreated cells, EGFR had a median *f_c_*value of 0.02 (Fig. 2A, white), consistent with it being primarily monomeric as reported by our lab previously(58, 59). We added an EGFR ligand, EGF, and observed a median *f_c_*value of 0.24 (Fig. 2A, gray), consistent with ligand-induced multimerization(59). The addition of EGF ligand to EGFR-also led to receptor internalization, which can be seen in the epifluorescence images (Fig. 2C). PIE-FCCS data was then collected after increasing the level of PIP_2_ by treating the cells with a PLC inhibitor (U73122, 5µM) for at least 15 minutes. The median *f_c_*value with the inhibitor was 0.10, indicating PIP_2_ induced dimerization of EGFR (Fig. 2A, blue). In contrast, decreasing PIP_2_ levels with a PLC activator (3M3FBS, 25 µM) resulted in a median *f_c_* value of 0.03 (Fig. 2A, green). Therefore, reduction of PIP_2_ did not significantly affect the multimerization nor diffusion of EGFR. The interpretation of the *f_c_* values is further supported by the diffusion coefficients of the receptors. The diffusion coefficients of EGFR decreased from 0.42 µm^2^/s to 0.15 µm^2^/s with EGF treatment (Fig. 2B, gray), and to 0.36 µm^2^/s after U73122 treatment (Fig. 2B, blue). There was no statistically significant change in the diffusion coefficient with the PLC activator (0.40 µm^2^/s) compared to EGFR in the resting cells (0.42 µm^2^/s).

**Figure 2.**
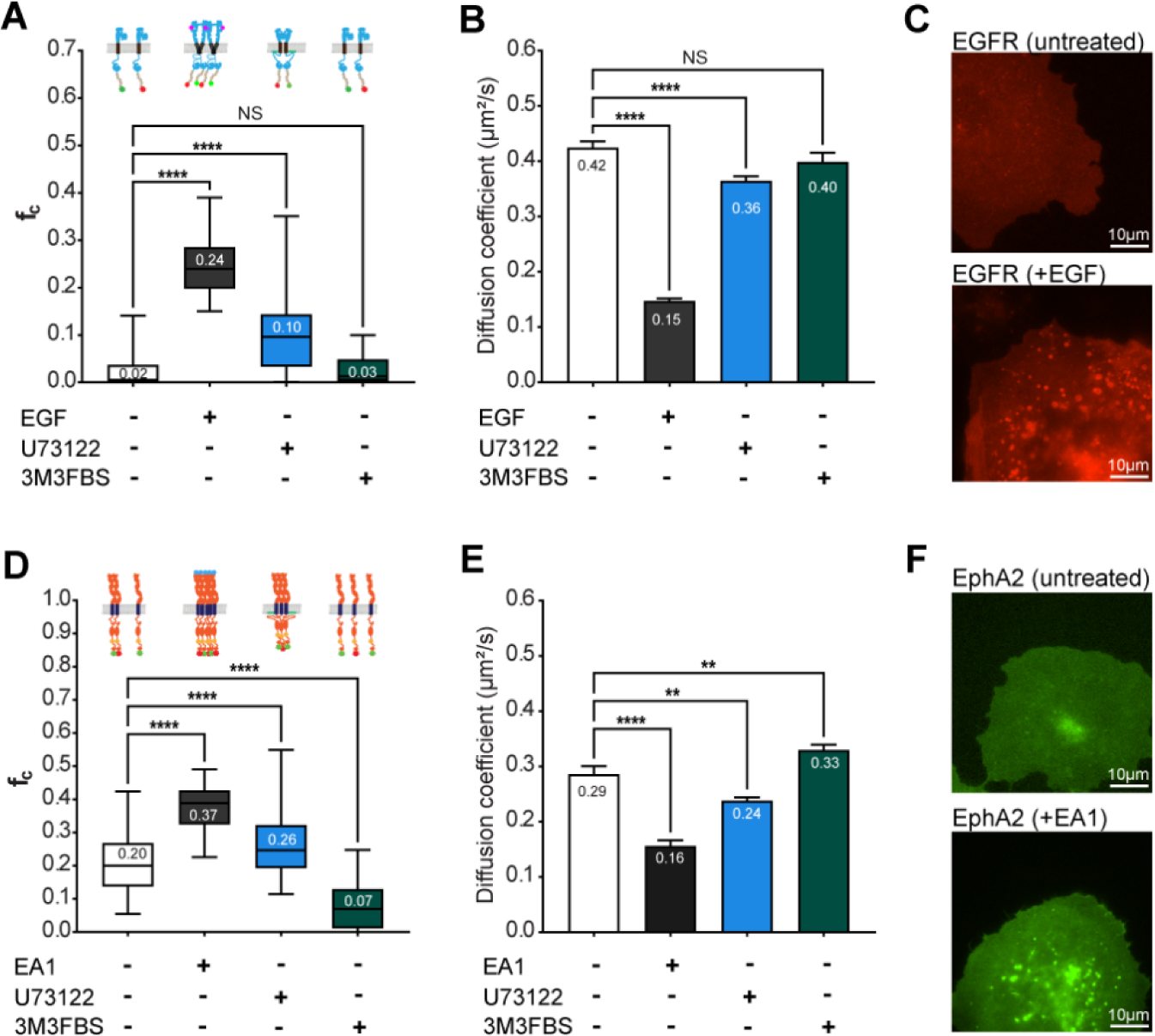
PIE-FCCS investigation of the homomeric interactions of EGFR and EphA2. **(A)** The self-assembly of EGFR was studied under various conditions. PIE-FCCS data was recorded and the resulting *f_c_*values are reported prior to treatment (white), after the addition of EGF (gray), after treatment with the PLC inhibitor U73122 (blue), and after treatment with the PLC activator 3M3FBS (green). **(B)** The diffusion coefficients for EGFR are shown for each treatment condition. **(C)** Epifluorescence images of EGFR expressing COS-7 cells without (top) and with (bottom) EGF ligand treatment. **(D)** The homo-multimerization state of EphA2 was recorded under the same treatments except the ligand treatment was with EA1. **(E)** The diffusion coefficients are shown for EphA2 under each treatment condition. **(F)** Epifluorescence images of EphA2 expressing COS-7 cells are shown without (top) and with (bottom) EA1 ligand treatment. In the box-and-whiskers plots, the whiskers indicate the maximum and minimum values; the box indicates the 25th-75th percentile, and the line is the median value (median value shown as text). We performed one-way ANOVA tests with uncorrected Fisher’s LSD post-hoc tests to obtain adjusted and individual P values (* = P<0.0367, ** = P<0.0034, **** = P<0.0001). Diffusion coefficient data are represented as mean values ± SEM.

To determine the effect of PIP_2_ levels on EphA2 homo-multimerization, we expressed EphA2-GFP and EphA2-mCh in Cos-7 cells. PIE-FCCS data were collected with and without ephrinA1 (EA1) or PLC regulators as described above. Prior to the ligand or drug treatment, the median *f_c_*value was 0.20, indicating substantial homo-multimerization (Fig. 2D, white) as we have reported recently(60). The *f_c_* values for EphA2 receptors without ligand were larger than those for dimer controls (Fig. 1C) and more comparable to those for multimer controls(47). We next added EA1, an EphA2-specific ligand, and observed a median *f_c_* value of 0.37 (Fig. 2D, gray), indicative of ligand-induced multimerization. The EphA2 diffusion coefficient decreased from 0.29 µm^2^/s (Fig. 2E, white) to 0.16 µm^2^/s (Fig. 2E, gray) with EA1 ligand treatment, which supports our median *f_c_* value interpretations. The addition of the PLC inhibitor (U73122, 5µM) resulted in a larger median *f_c_* = 0.27 (Fig. 2D, blue) compared to resting cells. Similar to the EGFR results above, we interpret this as PIP_2_-induced multimerization of EphA2. We next treated cells with a PLC activator (3M3FBS, 25µM), which resulted in a reduced median *f_c_* = 0.07. This result indicates that lower PIP_2_ reduced the ligand-independent multimerization of EphA2. The diffusion coefficients support these interpretations. The PLC inhibitor reduces protein mobility from 0.29 µm^2^/s (Fig. 2E, white) to 0.24 µm^2^/s (Fig. 2E, blue), and the PLC activator enhanced it to 0.33 µm^2^/s (Fig. 2E, green). Together, these results demonstrate that PIP_2_ induces EGFR homodimerization and EphA2 homomultimerization.

### PIP_2_ and EGF positively regulate EGFR-EPHA2 hetero-multimerization

We next tested the effect of PIP_2_ on the interaction between the EGFR and EphA2 receptors. Cos-7 cells were transfected with both receptors (EGFR-mCH and EphA2-eGFP) and PIE-FCCS measurements were performed as described above. During data collection, cells were selected with a nearly equal expression of both proteins. Under resting cell conditions, no statistically significant cross-correlation was observed between EGFR and EphA2, as indicated by the median *f_c_* value of 0.04 (Fig. 3A, white). We next incubated cells with 500 ng/ml of EGF or 500 ng/ml of EA1. PIE-FCCS data were collected between 15 and 90 minutes after ligand stimulation. After EGF treatment, the median *f_c_* value increased from 0.04 to 0.10, indicating the increased formation of hetero-multimers. This increase in cross-correlation is consistent with previous reports that EGF can enhance the association of EGFR and EphA2(30). Combined with our observations in the previous section, we conclude that the hetero-multimers consisted of EGFR dimers and EphA2 multimers, which could have a size range of 4 to 8 proteins. This interpretation is supported by the diffusion coefficient data in which we observe that the mobility of EGFR decreases significantly from 0.37 μm^2^/s to 0.15 μm^2^/s, while the diffusion coefficient of EphA2 reduced from 0.25 μm^2^/s to 0.20 μm^2^/s (Fig. 3 B & C). We next added the EphA2-specific ligand, EA1. In the presence of EA1, the median *f_c_* value was 0.02 (Fig. 3A, orange), statistically indistinguishable from the untreated cells, supporting that EA1 does not lead to hetero-multimerization between EGFR and EphA2. The reduction in the diffusion coefficient of the EphA2 receptor from 0.25 µm^2^/s to 0.16 µm^2^/s is explained by EA1-induced homo-multimerization of EphA2 (Fig. 3C).

**Figure 3.**
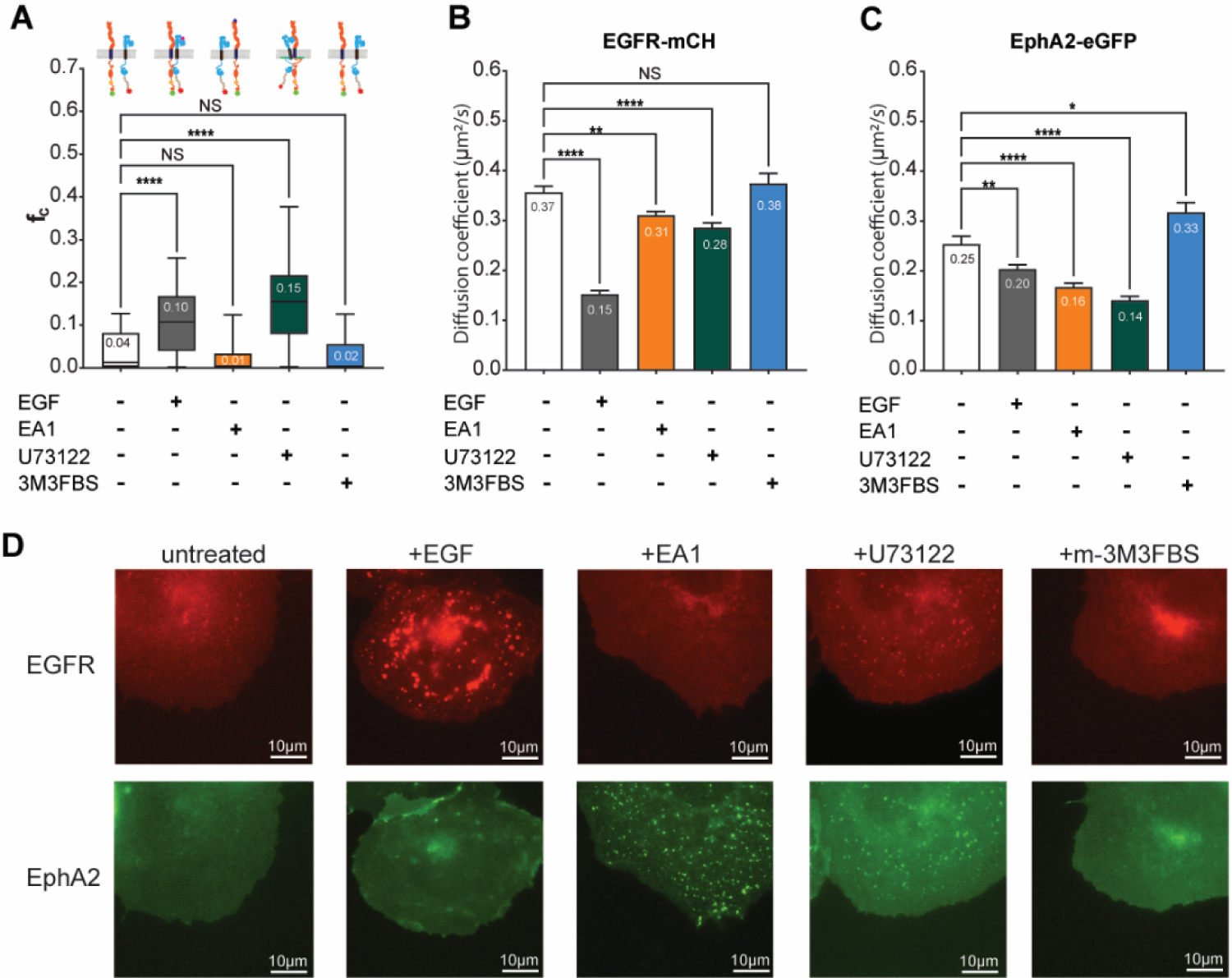
Hetero-multimerization of EGFR and EphA2 in the presence of ligands and PLC drugs. **(A)** PIE-FCCS data for EGFR-mCH and EphA2-eGFP co-expressed in COS-7 cells. Data were recorded prior to treatment (white), after the addition of EGF (gray) or EA1 (orange), and after treatment with the PLC inhibitor U73122 (blue), or with the PLC activator 3M3FBS (green). **(B)** Diffusion coefficients of EGFR-mCH after treatment with ligands and PLC-regulating drugs. **(C)** Diffusion coefficients of EphA2-GFP after treatment with ligands and PLC-regulating drugs. **(D)** Representative epifluorescence images from each set of experiments. One-way ANOVA tests with uncorrected Fisher’s LSD post-hoc tests shows the individual P values (* = P<0.0259, ** = P<0.003, **** = P<0.0001). Diffusion coefficient data are represented as mean values ± SEM.

We next tested the effect of the PLC inhibitor, and observed a significant change in the median *f_c_* value, from 0.04 to 0.15, indicative of heteromultimerization (Fig. 3A, green). The average diffusion coefficient of EGFR decreased from 0.37 µm^2^/s to 0.28 µm^2^/s. The mobility of EphA2 receptors also decreased from 0.25 to 0.14 µm^2^/s. The median *f_c_* value remained unchanged after adding the PLC activator compared to resting cells, as shown in Fig. 3A (blue). The molecular diffusion of EGFR indicated no change in protein mobility (Fig. 3B, blue), consistent with the *f_c_* values. The mobility of the EphA2 receptor increased after treatment with the PLC inhibitor, consistent with the disruption of homo-multimerization explained in the previous section. Lifetime analysis of eGFP shows a slight reduction in lifetime with ligand (EGF) and PLC inhibitor, while mCherry slightly increases. No statistically significant changes were observed with the EA1 ligand and PLC activator. The combined effect of ligand and PLC protein inhibitor (U73122) shows a slight increase in the median *f_c_*value of EGFR-EphA2 hetero-multimer.

### PIP_2_ -induced EGFR and EphA2 hetero-multimers are inactive

Once we determined how PLC drugs affected EGFR and EphA2 homo- and hetero-multimerization, we investigated the activity of the multimers. Phosphorylation of key tyrosine residues is commonly used as an indicator that RTKs are in their active conformation. Additionally, EphA2 S897 has been shown to be important for heterodimer formation and activity(30). We therefore utilized western blotting to determine if these phosphorylation sites were affected by the PLC drugs (Fig. 4). We began by treating A375 cells with EA1 and EGF to confirm the normal activation of endogenous proteins in our system. EGF caused a marked increase in EGFR pY1068, showing that EGFR was successfully activated. EGF treatment also caused an increase in EphA2 pS897 (Fig. 4C) correlating with EGFR-EphA2 heterodimerization. As expected, EA1 induced a strong increase in EphA2 pY588. We chose a 15-minute treatment time to minimize the amount of EphA2 endocytosis and lysosomal degradation resultant from activation. However, there was a small reduction in total EphA2 levels (Fig. 4D) due to degradation(61). Together, these results demonstrate the successful activation of EGFR and EphA2 homo- and hetero-multimers by their respective ligands.

**Figure 4.**
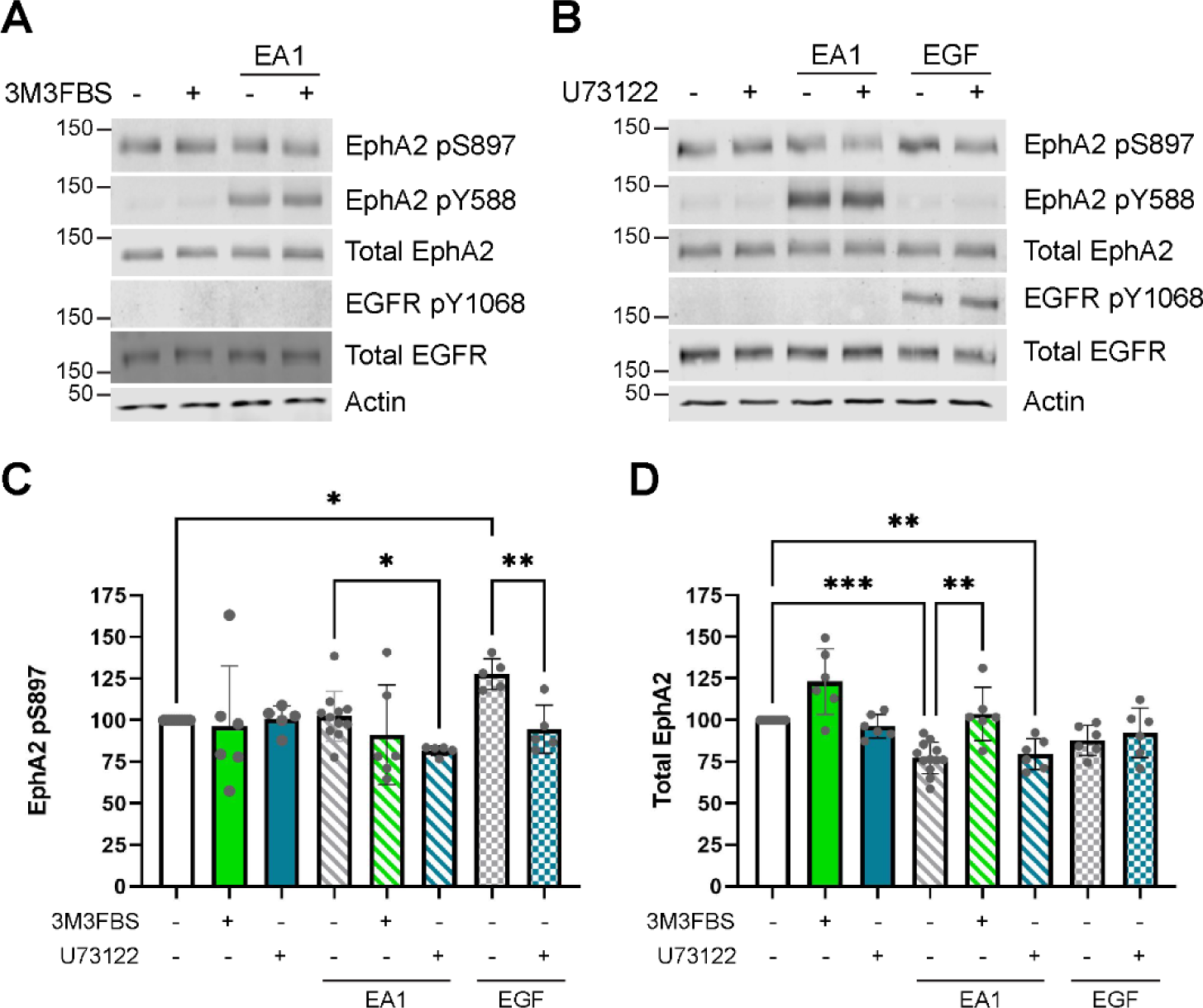
EGFR and EphA2 phosphorylation changes after treatment with ligand and PLC drugs. **(A),** Representative western blots after treatment with EA1 (15 min, 500 ng/mL), 3M3FBS (45 min, 25 µM), or 3M3FBS followed by EA1. **(B),** Representative western blots after treatment with U73122 (15 min, 5 µM), EA1 (as above), EGF (15 min, 100 ng/mL), or U73122 followed by each ligand. **(C)** Quantification of EphA2 pS897 levels in all conditions normalized to the corresponding total EphA2 bands. N = 6-12. Statistical analysis was performed using a Kruskal Wallis test -H(7) = 19.45, *p* = 0.007-with a Mann Whitney U test for comparisons between groups. Significance values were adjusted by the Bonferroni correction for multiple tests. **(D),** Quantification of total EphA2 levels in all conditions normalized to actin. N = 6-12. Statistical analysis was performed using a Kruskal Wallis test -H(7) = 38.44, *p* = 3.0 x 10^-6^ - with a Mann Whitney U test for comparisons between groups. Significance values were adjusted by the Bonferroni correction for multiple tests.

To determine the effects of PIP_2_ on EGFR and EphA2 phosphorylation, we treated A375 cells with 5 µM U73122 for 15 min or 25 µM 3M3FBS for 45 min. Neither up- nor down-regulation of PIP_2_ caused phosphorylation changes for EGFR pY1068 or EphA2 pY588, indicating that PIP_2_ changes do not trigger signaling associated to ligand-dependent activation. The PIE-FCCS data above show that PIP_2_ promotes self-assembly of each receptor (Fig. 2A, 2D), however, the phosphorylation data indicate that these are inactive multimers. We were intrigued to find that a similar phenomenon occurred for EphA2 pS897: regulation of PIP_2_ via PLC drugs had no effect on the level of EphA2 pS897 (Fig. 4C). This demonstrates that the PIP_2_-induced hetero-multimers do not affect the availability of EphA2 as an Akt substrate(62). Altogether, the functional data support the conclusion that PIP2 induced EGFR-EphA2 hetero-multimers trap the proteins in an inactive state.

We also investigated if ligand activation of EGFR and EphA2 was affected by PIP_2_ levels. We treated A375 cells first with the PLC modifiers, followed with EGF or EA1 treatment. Treatment with EA1 after downregulation of PIP_2_ had no effect on the levels of EphA2 pY588. This suggests that EA1 can induce activation of EphA2 regardless of the initial oligomer size. Intriguingly, EA1 did not cause a decrease in total EphA2 in cells with decreased PIP_2_ (Fig. 4D). Though activation occurs normally, this suggests that there may be a difference in the ability of activated EphA2 to be degraded after ligand treatment. EGF treatment after downregulation of PIP2 was not tested, as 3M3FBS had no effect on EGFR homodimerization nor EGFR-EphA2 heterodimerization (Fig. 2 & 3).

EA1 treatment of cells containing higher PIP_2_ caused EphA2 pY588 to increase to the same level as cells with basal amounts of PIP_2_. This result suggests that PIP_2_-mediated EphA2 homomultimers can still convert to active multimers successfully. Upregulation of PIP_2_ prior to EA1 treatment decreased pS897 levels (Fig. 4C) similar to EGF treatment of cells containing upregulated PIP_2_. There was no PIP_2_-induced change in EGFR pY1068, suggesting successful activation of EGFR homomultimers, similar to that of EphA2 activation. However, when PIP_2_ levels were increased prior to EGF treatment, the increase in EphA2 pS897 characteristic of EGF treatment alone did not occur (Fig. 4C). Because EphA2 pS897 was decreased by both EA1 and EGF in high PIP_2_ cells, we hypothesize that this change is due to conformational differences resulting from heterodimerization.

### AlphaFold Multimer predicts a conformational change in the EGFR-EphA2 heterodimer

Based on our data, we conclude that EGFR-EphA2 hetero-multimers have two conformations: an inactive conformation that is not EphA2 S897 phosphorylated and an active conformation where is S897 phosphorylated. Our results additionally indicate that PIP_2_ stabilizes the inactive complex. There are no structures for EGFR-EphA2 heterodimers. However, recent advances in artificial intelligence allow us to obtain robust predictions of protein complexes. AlphaFold Multimer (AlFoM) is an extension of AlphaFold2 developed to predict protein-protein complexes(63). We applied AlFoM to predict the structure of the inactive and active EGFR-EphA2 heterodimer. We performed AlFoM simulations of the EGFR-EphA2 intracellular domains (ICDs), which have been proposed to interact (64). The results show asymmetric kinase dimerization (Fig. 5A) similar to that observed for the active state of EGFR homodimers(65). To benchmark the predicted structures, they were overlayed with crystal structures for EGFR (PDB: 2GS6) or EphA2 (PBD: 7KJA) (64, 66), and the RMSD was 2.9 Å and 8.7 Å, respectively. The larger RMSD for EphA2 can be attributed to the change in the conformation of the SAM domain, but SAM domain movement is both expected and necessary for EphA2 activation. Therefore, to obtain a more accurate RMSD that reflects the structural similarities between the predicted and experimental structures while removing the confounding factor of the SAM domain orientational change, we split the EphA2 ICD into two pieces and repeated the RMSD calculation. EphA2 residues 559 to 875 -which cover the kinase domain-overlay with our predicted structure with an RMSD of 1.8 Å, while SAM domain residues 904 to 976 overlay with an RMSD of only 0.8 Å, suggesting that the predicted structures are supported by experimentally determined structures. We additionally plotted the IDDT values per residue, showing a relatively high level of confidence in these predictions over the structured portions of the proteins.

**Figure 5.**
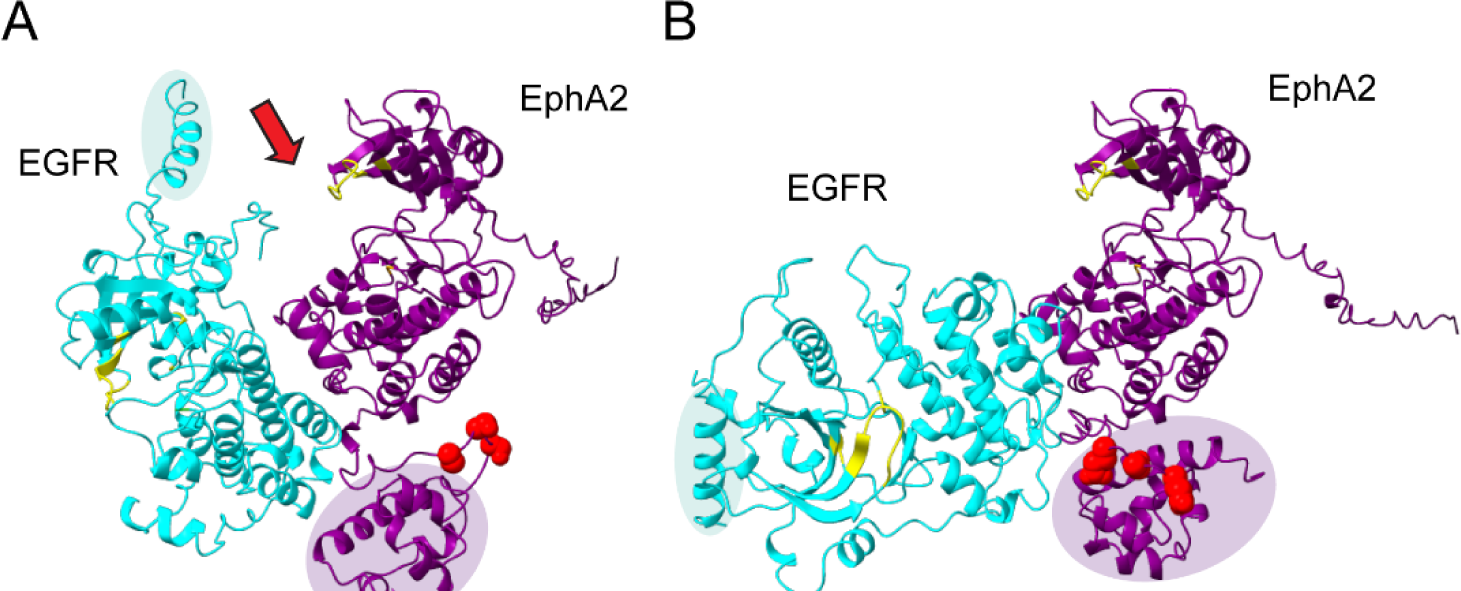
AlphaFold Multimer model of the heterodimer of EGFR residues 669-1210 (cyan) and EphA2 residues 559-976 (purple). Predictions are shown for EphA2 WT **(A)**, or for the S897E/S901E mutant **(B)** representative of phosphorylation at serine residues (red bubbles). Yellow residues indicate the active site of each kinase region. The purple halo encompasses the EphA2 SAM domain while the cyan halo shows the EGFR JM domain. The red arrow indicates the EphA2 kinase active site which is sterically blocked by the EGFR kinase domain in A, but not in B.

It was previously reported that EphA2 phosphorylation at S897 and S901 causes a configurational change of the SAM domain that allows a change in the function of the EphA2 homodimer(64). Additionally, our western blot data show that S897 is also an essential factor in signaling by the EGFR-EphA2 heterodimer. We therefore repeated the AlFoM prediction with a previously validated phosphomimetic mutant of EphA2 in which S897 and S901 have been mutated to glutamic acid (EphA2_S897E/S901E_, Fig. 5B)(64). In this prediction, the RMSD of EGFR with its crystal structure provided an RMSD of 2.9 Å. The two-part EphA2 comparison was repeated, and the kinase domain had an RMSD of 1.8 Å while for the SAM domain was only 0.7 Å, showing excellent agreement between the crystal structure and the AlFoM model.

We were intrigued to find that pS897-mimetic mutations caused a large difference in the orientation of EGFR with respect to wild-type EphA2. Specifically, when EGFR interacts with EphA2_WT_, EGFR is positioned such that the active site of the EphA2 kinase domain (Fig. 5, yellow) is partially obstructed, while EGFR interacting with EphA2_S897E/S901E_ results in a more open kinase active site. We further noticed that the EGFR JM helix is positioned differently in each structure. The positively charged residues in the EGFR JM strongly interact with negatively charged PIP_2_, and this interaction regulates EGFR homodimer stability and activity(67-70). We therefore hypothesized that this interaction might regulate the conformation of the EGFR-EphA2 heterodimer as well. Based on the AlFoM model results, we hypothesize that when PIP_2_ levels are increased, the JM is sequestered at the membrane, disallowing the flexibility required for EGFR to shift into its more open and active conformation. Further experimental investigation into this model is required, but these structural predictions offer interesting possibilities for both EGFR-EphA2 dimer conformations and the effects of PIP_2_.

## Discussion

Though research over several decades has characterized RTK homodimer states, there is still much that is unknown about hetero-multimerization, including the conditions under which each multimer state is present. Our data indicate that EGFR-EphA2 heterocomplexes are not present under basal conditions or after EA1 treatment. Only in the presence of EGF do we detect hetero-multimerization in a timeframe of 15-90 minutes (Fig. 3). Previous reports of EGFR-EphA2 interaction conditions are contradictory. One study reported that endogenous EphA2 and EGFR showed increased co-immunoprecipitation (Co-IP) after EGF treatment, but only after 24 hours(25). Co-IP only occurred under basal conditions in cell lines where EGFR was constitutively activated/phosphorylated, which suggests that EGFR activation is necessary for heterodimer formation(25). A more recent study that utilized over-expressed proteins reported an increase in Co-IP in as little as 5-30 min after EGF treatment(30). While it appears that the time at which heterodimerization occurs could be dependent on cell line, these studies agree with our findings that activation of EGFR by EGF induces formation of EGFR-EphA2 hetero-multimers. However, another recent study reported the opposite: FRET showed that heterodimers are strong under basal conditions and are decreased by EGF treatment(71). These experiments were predominantly performed with truncated versions of the receptors that do not contain the ICDs, suggesting that the ICDs regulate the response to EGF. The importance of the ICDs is supported by our findings in the sense that PIP_2_ lipids localize to the inner leaflet of the plasma membrane and thus are expected to interact directly with the ICDs. Future work will be essential to determine how protein mutations, expression levels, different cell lines, and variations in signaling networks affect ligand-mediated EGFR-EphA2 heterodimer formation.

Our data show that PIP_2_ is an important factor in membrane receptor assembly. Because EGFR is one of the most well-studied RTKs, a notable effort has been undertaken to understand how PIP_2_ affects EGFR homodimers. Both computational and experimental studies have shown that PIP_2_ binds to the juxtamembrane (JM) segment via electrostatic interactions to stabilize the active conformation of EGFR homo-multimers(68, 69). Additionally, western blot studies of EGFR phosphorylation suggest that PIP_2_ facilitates the activation of EGFR by EGF(70). However, while we observed homodimer stabilization under high PIP_2_ conditions (Fig. 2), our western blot data did not support the hypothesis that PIP_2_ levels alone could drive ligand-independent EGFR phosphorylation. One possibility for this discrepancy is the difference in our methods used to disrupt PIP_2_. Michailidis *et al*. worked in oocytes treated with wortmannin, a PI4K inhibitor that decreases PIP_2_ levels by blocking lipid synthesis, rather than affecting degradation like U73122 and 3M3FBS. Additionally, PIP_2_ changes do not have an effect on EGFR phosphorylation in HeLa cells(70), indicating the effect might be cell-line dependent. It is important to continue to study why PIP_2_ level changes have an effect in some cells but not in others, as this effort will result in a better understanding of the molecular mechanisms of RTK heterodimerization.

A large-scale MD study suggested that PIP_2_ binding to the JM region is conserved across RTKs(33). Hedger and co-workers found that the conserved, positively-charged JM residues of all 58 known RTKs bind to and induce clustering of anionic lipids, including PIP_2_. The Barrera lab recently showed that PIP_2_ specifically promotes dimerization of a construct that contains the TM domain and a short JM span of EphA2. Interestingly, this effect was specific, as it only occurred in the ligand-independent dimer conformation(38). However, prior to this study, it was not known if PIP_2_ played a role in the multimerization of full-length EphA2. Here, we report that PIP_2_ promotes the formation of EphA2 homo-multimers (Fig. 2). Intriguingly, tyrosine autophosphorylation does not occur in these multimers, indicating that they are inactive. This result agrees with Stefanski *et al*.’s hypothesis that PIP_2_ preferentially promotes ligand-independent multimerization. This is particularly interesting, as unliganded EphA2 oligomers promote tumorigenesis(62, 72). Based on our data, we propose that tumors with elevated levels of PIP2 might suffer worse prognosis than those with lower levels. If this idea were correct, pharmacological regulation of PIP2 levels might be clinically beneficial to control EphA2 mediated cancers. Future work is necessary to test this hypothesis.

Since PIP_2_ promotes homodimerization of both EGFR and EphA2, it would be plausible to assume that it would reduce heteromultimerization simply by the laws of mass action. However, we were surprised to find that PIP_2_ also promotes heteromultimer formation (Fig. 3). Our study sets the stage for future work to determine how other anionic lipids like PS, or cholesterol, might regulate the heterodimerization of other RTKs, as well as the switch between homo- and hetero-dimerization. The phosphorylation data presented above suggest that PIP_2_ promotes an inactive form of the EGFR-EphA2 heteromultimer (Fig. 4C). Therefore, our data puts forward a mechanism by which there may be more than one EGFR-EphA2 functional state. Since RTKs including EphA2 and EGFR have more than one homodimer conformation relating to active and inactive states(3, 62, 73, 74), it is not unreasonable to suggest that this extrapolates to hetero-multimers.

In accordance with the finding presented here, we propose the following model outlined in Fig. 6. Under basal conditions, EGFR and EphA2 do not bind, and only form heterocomplexes in response to EGF activation. The EGF-induced hetero-multimer is phosphorylated at EGFR Y1068, corresponding to EGFR kinase activity, and EphA2 S897, corresponding to kinase activation and the availability of EphA2 as a substrate for serine/threonine kinases(62). The AlFoM predictions provide some insight to the conformation of inactive and active heterodimers (Fig. 5). Low cellular PIP_2_ allows the JM to move freely such that the kinase domains adopt an active conformation where the active pockets are free to perform phosphorylation. In contrast, increased levels of PIP_2_ cause the EGFR JM to bind tightly to the membrane. Such conformation would promote an inactive heterodimer conformation, where the EphA2 active pocket is sterically blocked by the EGFR kinase domain, hindering EphA2 phosphorylation activity. It is also possible that PIP_2_ interactions work in concert with the rotomeric state of the helical JM domain of EGFR, which is important for ligand-induced conformational coupling across the membrane(75). These mechanistic details will require additional structural studies, but our model provides an important conceptual framework that will help guide future experiments. More work is needed to establish the mechanism for the conformational transition from an inactive to active heterodimer, as well as how PIP_2_ affects EGFR-EphA2 heterodimers.

**Figure 6.**
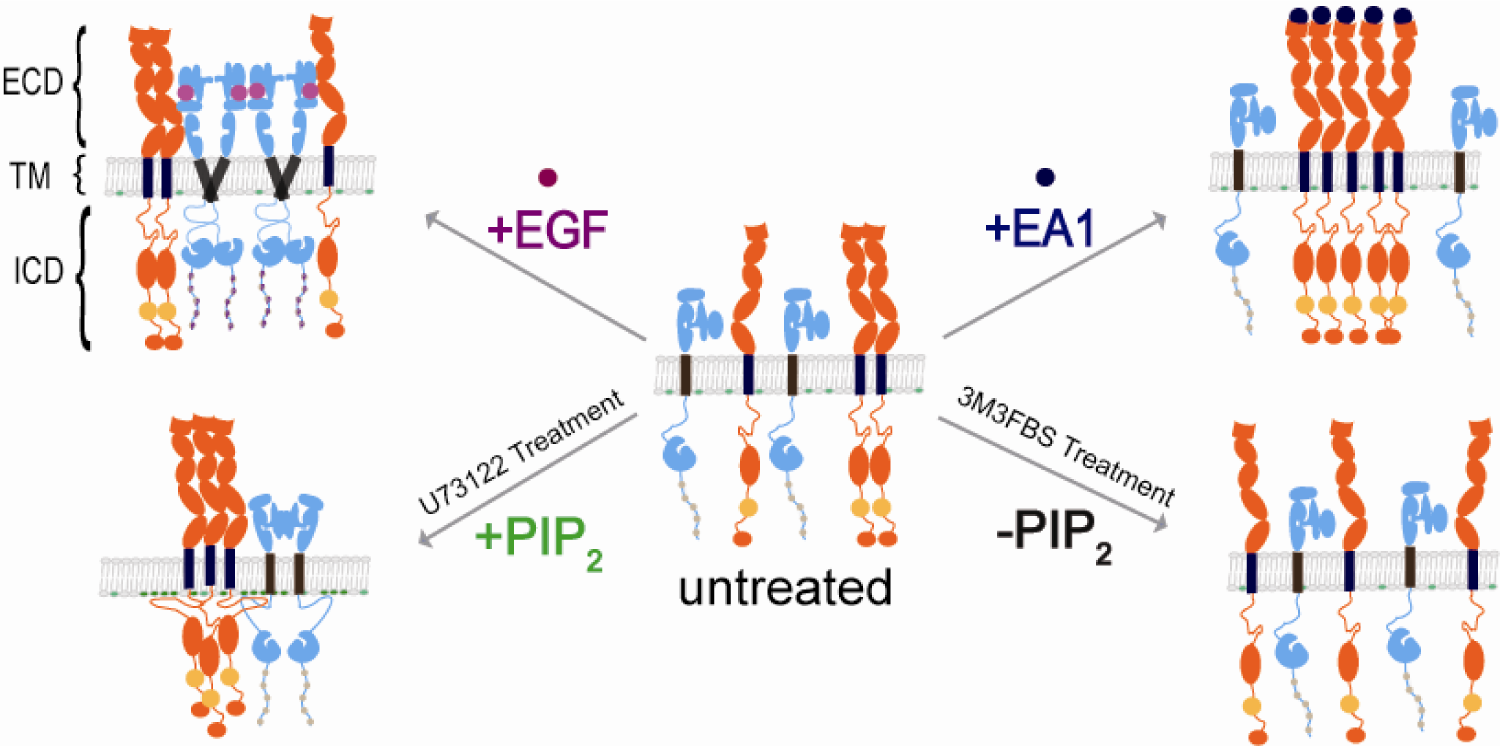
Model for EGFR-EphA2 heterodimerization. When PIP2 levels are normal, EGFR (blue) and EphA2 (orange) do not interact in the absence of ligand. When EGF is present, active heterodimer forms in which EGFR Y1068 and EphA2 S897 are phosphorylated. The active conformation of the ICDs allows a more open EphA2 kinase domain with the EGFR JM free to move away from the membrane. Under increased PIP_2_ conditions, PIP_2_ promotes inactive heterodimer formation. The inactive dimer is likely stabilized by the EGFR JM electrostatically interacting with the membrane and the EGFR ICD is pulled towards the membrane to close the EphA2 kinase domain.

## Materials and Methods

### Plasmids and Cloning

EGFR was subcloned to eGFP-N1 and mCherry-N1 vectors by XhoI and AgeI digest as described previously(76). Human EphA2 cDNA (AA 1-971 based on NM_004431.4) with C-terminal GFP and mCherry tag were purchased from Origene in gateway donor vectors as described previously(60). PH-PLCD1-GFP was a gift from Tamas Balla (Addgene plasmid # 51407; http://n2t.net/addgene:51407; RRID:Addgene_51407) (77).

### Cell culture and ligand stimulation

For the culture and expression of the receptor proteins, the Cos-7 cells were transiently transfected using standard methods. This study used Dulbecco’s modified Eagle’s medium supplemented with 10% fetal bovine serum and 1% penicillin-streptomycin. Transfection of the plasmids was carried out with 70 to 80% confluent cells in 35 mm glass-bottom MatTek dishes. The cells were transiently transfected approximately 20 hours before the data collection with the protein of interest using Lipofectamine 2000 reagent (Thermo Fisher Scientific). A total of 2.5 μg DNA (1:1) ratio of mCherry and eGFP-tagged plasmids was used to express both fluorescent tags evenly and acquire a local density of 100–1500 receptors/μm^2^. Before measuring FCCS, the media was changed to Opti-MEM I Reduced Serum Medium (Thermo Fisher Scientific) without phenol red. For each complex, measurement was taken for both the ligand-free and ligand-stimulated state, using either recombinant human EGF (Sigma-Aldrich, St. Louis, MO) or Ephrin A1 as ligand. In order to stimulate receptor-expressing cells, a stock solution (20 µg/ml) was diluted to 500 ng/ml in Opti-medium (imaging media) and added approximately 15 minutes prior to data collection. Data was collected for a maximum of one hour following stimulation.

We have previously published control constructs useful in interpreting *f_c_* values for monomers, dimers, and higher-order oligomers. Moreover, these control constructs indicate that the fluorescent proteins themselves do not promote or inhibit dimerization states of the proteins. According to a homotypic interaction, the *f_c_* values below 0.09 indicate monomers, 0.09-0.17 indicate dimers, and above 0.17 indicate higher-order oligomers.

Cells were incubated with different concentrations of PLC regulators to determine the optimum concentration. The PLC inhibitor (U73122) was incubated for 15 minutes at 1.0 - 5.0 µM concentration prior to data collection. Concentration ranges of 1.0 to 4.0 did not show any significant changes; however, concentration ranges of 5.0 did show a significant change. After 15 min of PLC activator (m-3M3FBS) stimulation, EFGR/EphA2 did not show a detectable *f_c_*value change with 5 µM, 10 µM, 15 µM, and 20 µM concentration. However, the further experiments with 25 µM concentration and 45 minutes incubation, the FC value of cross-correlation changed from 0.20 to 0.04, indicating monomeric forms of EphA2 receptors.

### PIE-FCCS instrumentation, data collection, and analysis

The time-time autocorrelation function is employed for single color FCS to analyze intensity fluctuations. The results were plotted on a semi-log axis to better view the time scale. In principle, ACF amplitude is inversely proportional to the average number of molecules in the observation area. In FCCS, two fluorescent probes are used to analyze the emission independently of one another using two separate spectrally distinct probes. As a result, both populations have a corresponding ACF, and molecular density can be determined independently. With PIE-FCCS, two laser pulses are phase-delayed calculating the exact arrival time of each laser pulse’s emission photons.

The FCCS data were recorded using pulsed interleaved excitation and time-correlated single-photon detection with a custom inverted microscope setup. A supercontinuum pulsed laser (9.2 MHz repetition rate, SuperK NKT Photonics, Birker*d, Denmark) was split into two beams of 488 and 561 nm using a series of filter and mirror combinations. In order to achieve PIE, the beams are directed through separate optical fibers of varying lengths, causing a delay in arrival time between them. This eliminates spectral crosstalk between the detectors. Before entering the microscope, the beams were overlapped by a dichroic beam splitter (LM01-503-25, Semrock) and a customized filter block (zt488/561rpc, zet488/561m, Chroma Technology). The overlapping beams of light were focused by the objective (×100 TIRF) to a limited diffraction spot on the peripheral membrane of a Cos-7 cell expressing the receptor constructs. In time-tagged time-resolved mode, photons were detected by individual avalanche photodiodes (Micro-Photon Devices). In order to verify the alignment of the system, including the overlap of a confocal volume, a short fluorescently tagged DNA fragment was used. Prior to the experimental samples, positive and negative controls (Fig. 1D-E) were tested to compare the fit parameters.

The excitation beams were focused on the peripheral membranes to measure only membrane-bound receptors. Only the cell’s flat, peripheral membrane area was scanned to prevent fluorescence from cytosolic organelles or vesicles. The data collection was performed on one area of the membrane in each sample. Each area was assessed six times, with each acquisition lasting 10 seconds. MATLAB scripts were used to calculate each sample’s auto- and cross-correlation curves. As described in the previous work, we fit a single component, the 2D diffusion model, to the averaged curves of six consecutive acquisitions per area.

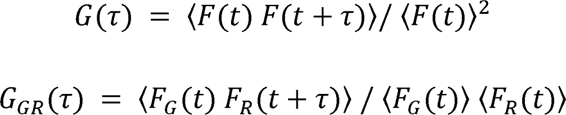

The intensity fluctuations were gated using PIE prior to cross-correlation and autocorrelation analyses. The intensity at time F(t) was compared to the intensity at a later time F(t + ⍰) in an autocorrelation analysis. As a function of time, self-similarity allowed for the interpretation of quantitative information, such as diffusion and particle number. In Equation 1, the intensity fluctuations were divided into 10-second bins and normalized to the square of the average intensity. The autocorrelation functions were fit to the following model for two-dimensional diffusion in the membrane that accounts for triplet relaxation and dark state dynamics.

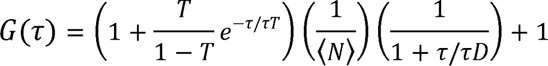

The diffusion coefficient of the sample has been calculated by utilizing the standard formula given below.

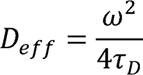

Finally, the cross-correlation function (fraction of correlation) was by using the autocorrelation function of both fluorophores.

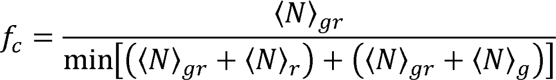

### Control samples for fraction of correlation (*f_c_*) calculation

We applied two samples for the calculation of fc value of membrane proteins. The first, a monomer, is the SRC derived peptide fused directly to the fluorescent protein. The second, a dimer control, is a lipid-anchored peptide derived from SRC fused to the GCN4 (SRC-GCN4) α-helical dimerization motif and a C-terminal attached fluorescent protein i.e. eGFP or mCherry. The SRC and SRC-GCN4 constructs were used as negative and positive controls, respectively, for cross-correlation in the experiment. The absence of any cross-correlation in SRC indicates that PIE has successfully eliminated any artifacts that could have caused inaccurate cross-correlation readings, and that the fluorescent proteins themselves did not play a role in promoting dimerization. The data from SRC-GCN4 demonstrate the maximum limit for a system that is strongly dimerized, which can be attributed to protein dark states and the existence of dimers having the same fluorescent protein tags. Laser powers were set to 0.350 μW and 0.80 μW for the 488 nm and 561 nm lasers, respectively.

To calibrate the highest fluorescence value of a sample, we utilized double-labeled DNA strands. These strands were also used as DNA standards to calibrate confocal volumes. A 40-nucleotide sequence with <50% G-C content [ACA AGC TGG AGT ACA ACT ACA ACA GCC ACA ACG TCT ATA T] was labeled with carboxy-tetramethylrhodamine (TAMRA) at the 5’ end and 6-carboxyfluorescein (FAM) at the 3’ end (Integrated DNA Technologies). An excess of the unlabeled strand was used to anneal the synthesized complementary strand pair as per the supplier’s protocols. The double-stranded DNA, labeled with both TAMRA and FAM (TAMRA-40-FAM), was diluted to a final concentration of 100 nM using 10 mM TE buffer. For the 3D sample data collection, laser powers were set to 7 μW and 7 μW for the 488 nm and 561 nm lasers, respectively.

### Live-cell microscopy of PLC**δ**_1_PH-GFP

Live cell imaging was performed to verify the effect of the PLC drugs in an assay adapted from Balla et al(57). To prepare cells for imaging, A375 cells were plated at 60% confluency on a #1.5 glass-bottom 8-well chamber slide. The glass was coated in fibronectin (10 µg/mL) for 30 min at 37°C prior to plating, and cells were allowed to adhere for 24 hr. Cells were then transfected in OptiMEM media with PLCδ_1_PH-GFP using a 3:1 ratio of polycation polyethylenimine (PEI) to DNA with 0.2 µg of DNA per well and allowed to incubate overnight. Cells were washed using PBS, starved of serum for 4 hours, and stained with Hoechst 33342 for 20 min to allow visualization of the nucleus. Cells were finally placed in FluoroBrite media for optimal imaging.

Timelapse imaging of cells were performed using an inverted Zeiss LSM 900/Airyscan laser scanning confocal microscope (ZEISS) with a 63X oil immersion objective. For a given well, 3-6 spots containing multiple fluorescent cells were chosen. The well was then treated with 0.5% (v/v) DMSO, and the desired Z-plane was found using DefiniteFocus. For cells planned for U73122 treatment, images were taken every 5 min for 20 min, and for cells planned for 3M3FBS treatment, images were taken every 10 min for 50 min. Following the experiment, DMSO was replaced with either U73122 or 3M3FBS, the DefiniteFocus was re-established, and timelapse images were repeated as above.

Images were processed and quantified using the ImageJ software. Four representative membrane areas and four representative cytosolic areas were chosen. The mean pixel intensity for each section was determined, and the normalized ratio of membrane to cytosol intensity for each single cell was calculated using the following equation:

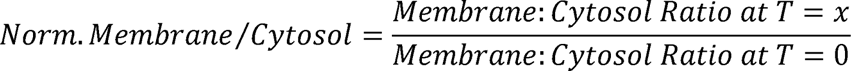

### Western Blot

A375 cells were plated to approximately 80% confluency in a 12-well culture dish and allowed to adhere for 24 hours. Cells were then starved of serum overnight. Treatments were made in serum-free DMEM, and cells were treated with 5 µM U73122 for 15 min or 25 µM 3M3FBS for 45 min then treated with 500 ng/mL EA1 or 100 ng/mL EGF for 15 min. Control cells were treated with equal amounts of DMSO as PIP_2_ modifying drugs. Once treatments were complete, cells were scraped from the plate and lysed in TEN-T lysis buffer (50 mM tris (pH 7.5), 100 mM sodium chloride, 1 mM EDTA, 1% triton X-100) containing protease and phosphatase inhibitors for 30 min on ice. Lysates were centrifuged at 13,000 x g for 10 min to remove insoluble material. 10 ng of protein was run on a 10% SDS-PAGE gel and transferred to 0.45 µm nitrocellulose. Membranes were blocked with 5% milk for total protein blots or 3% BSA for phospho-protein blots for 1 hr at room temperature and then incubated in primaries overnight at 4 °C. Secondaries were incubated 1 hr at room temperature and imaged for fluorescence on an Odyssey CLx Imaging System (LI-COR, Lincoln, Nebraska). Bands were quantified using Image Studio Lite. Phospho-protein bands were normalized to the corresponding total protein levels, while total protein bands were normalized to their corresponding actin loading control bands.

| Antibody | Catalog # | Solvent | Dilution | Host Species |
| --- | --- | --- | --- | --- |
| Total EGFR | 4267S, CST | 5% Milk in TBS | 1:1000 | Rabbit |
| pEGFR(Y1068) | 3777S, CST | 3% BSA in TBS | 1:1000 | Rabbit |
| Total EphA2 | 97535S, CST | 5% Milk | 1:1000 | Rabbit |
| pEphA2(Y588) | 12677, CST | 3% BSA | 1:1000 | Rabbit |
| pEphA2(S897) | 6347S, CST | 3% BSA | 1:1000 | Rabbit |
| Actin | 3700S, CST | 5% Milk | 1:5000 | Mouse |
| Anti-Mouse | 925-68020, LiCore | 5% Milk | 1:5000 | Goat |
| Anti-Rabbit | 926-32211, LiCore | 5% Milk | 1:5000 | Goat |

### AlphaFold Multimer

AlphaFold Multimer (56) was used to predict ICD dimerization of:

EGFR(R^669^RRHIVRKRTLRRLLQERELVEPLTPSGEAPNQALLRILKETEFKKIKVLGSGAFGTVYKGLWIPEGEKVKIPVAIKELREATSPKANKEILDEAYVMASVDNPHVCRLLGICLTSTVQLITQLMPFGCLLDYVREHKDNIGSQYLLNWCVQIAKGMNYLEDRRLVHRDLAARNVLVKTPQHVKITDFGLAKLLGAEEKEYHAEGGKVPIKWMALESILHRIYTHQSDVWSYGVTVWELMTFGSKPYDGIPASEISSILEKGERLPQPPICTIDVYMIMVKCWMIDADSRPKFRELIIEFSKMARDPQRYLVIQGDERMHLPSPTDSNFYRALMDEEDMDDVVDADEYLIPQQGFFSSPSTSRTPLLSSLSATSNNSTVACIDRNGLQSCPIKEDSFLQRYSSDPTGALTEDSIDDTFLPVPEYINQSVPKRPAGSVQNPVYHNQPLNPAPSRDPHYQDPHSTAVGNPEYLNTVQPTCVNSTFDSPAHWAQKGSHQISLDNPDYQQDFFPKEAKPNGIFKGSTAENAEYLRVAPQSSEFIGA^1210^)and

EphA2_WT_(H^559^RRRKNQRARQSPEDVYFSKSEQLKPLKTYVDPHTYEDPNQAVLKFTTEIHPSCVTRQKVIGAGEFGEVYKGMLKTSSGKKEVPVAIKTLKAGYTEKQRVDFLGEAGIMGQFSHHNIIRLEGVISKYKPMMIITEYMENGALDKFLREKDGEFSVLQLVGMLRGIAAGMKYLANMNYVHRDLAARNILVNSNLVCKVSDFGLSRVLEDDPEATYTTSGGKIPIRWTAPEAISYRKFTSASDVWSFGIVMWEVMTYGERPYWELSNHEVMKAINDGFRLPTPMDCPSAIYQLMMQCWQQERARRPKFADIVSILDKLIRAPDSLKTLADFDPRVSIRLPSTSGSEGVPFRTVSEWLESIKMQQYTEHFMAAGYTAIEKVVQMTNDDIKRIGVRLPGHQKRIAYSLLGLKDQVNTVGIPI^967^).

For EphA2_S897E/S901E,_ the two bolded S residues were replaced with E. The crystal structures of EGFR (PDB: 2GS6) (59) and EphA2 (PDB: 7KJA) (60) were obtained from https://www.rcsb.org/. For each simulation five models were provided, and the Best Model was used for further analysis. RMSDs were calculated using the ChimeraX matchmaking tool.

## Conflict of Interest

The authors declare no conflict of interests.

## Acknowledgements

We are thankful to Tim Simmons and Amit Joshi for experimental advice and the use of the LSM 900/Airyscan microscope.

## Funding

This work was supported by the National Science Foundation grant CHE-1753060, the Human Frontiers of Science Program No. RGP0059/2019, the American Lung Association Lung Cancer Discovery Award, LCD-1035035 (to AWS) and NIH grant R35GM140846 (to FNB).

## Author contributions

Conceptualization: AWS, PKS, FNB; Performed Experiments: PKS, JAR, RJS; Data Analysis: PKS, AWS, JAR, FNB; Writing original draft: PKS, AWS, JAR, FNB; Writing-Review and editing: AWS, FNB, PKS, JAR; Supervision and Project Administration: AWS, FNB; Funding Acquisition: AWS, FNB.

## Notes

### Competing Interest Statement

The authors have declared no competing interest.

